# Implicit reward-based motor learning

**DOI:** 10.1101/2023.06.27.546738

**Authors:** Nina M. van Mastrigt, Jonathan S. Tsay, Tianhe Wang, Guy Avraham, Sabrina J. Abram, Katinka van der Kooij, Jeroen B. J. Smeets, Richard B. Ivry

## Abstract

Binary feedback, providing information solely about task success or failure, can be sufficient to drive motor learning. While binary feedback can induce explicit adjustments in movement strategy, it remains unclear if this type of feedback also induce implicit learning. We examined this question in a center-out reaching task by gradually moving an invisible reward zone away from a visual target to a final rotation of 7.5° or 25° in a between-group design. Participants received binary feedback, indicating if the movement intersected the reward zone. By the end of the training, both groups modified their reach angle by about 95% of the rotation. We quantified implicit learning by measuring performance in a subsequent no-feedback aftereffect phase, in which participants were told to forgo any adopted movement strategies and reach directly to the visual target. The results showed a small, but robust (2-3°) aftereffect in both groups, highlighting that binary feedback elicits implicit learning. Notably, for both groups, reaches to two flanking generalization targets were biased in the same direction as the aftereffect. This pattern is at odds with the hypothesis that implicit learning is a form of use-dependent learning. Rather, the results suggest that binary feedback can be sufficient to recalibrate a sensorimotor map.

## Introduction

The execution of accurate movements relies on sensory feedback. Variants of sensorimotor adaptation experiments have been used to study the role of different forms of feedback on motor learning. In a typical visuomotor adaptation experiment, participants perform target-directed center-out reaching movements with feedback of the unseen hand limited to a visual cursor. To study learning, the position of the cursor is altered, resulting in a sensory prediction error, defined by the difference between the predicted and actual cursor position (Izawa & Shadmehr, 2011; Kim et al., 2018; Morehead et al., 2017; Shadmehr et al., 2010; Synofzik et al., 2008; Tseng et al., 2007). This directional error can drive different forms of learning. It can produce recalibration of a so-called sensorimotor map such that a subsequent movement to that target will be shifted in the direction opposite to the perturbed feedback, a process known as sensorimotor adaptation (Kim et al., 2021; Krakauer, 2009; Krakauer et al., 2019). It can also elicit explicit strategies to reduce the error; for example, the participant might aim away from the target (Bond & Taylor, 2015; Taylor et al., 2014).

Feedback can also be limited to binary information conveying success or failure. In reaching tasks, success can be defined by the hand intersecting an invisible reward zone. To elicit learning, the reward zone is displaced from the target. This might be done in an abrupt manner. For example, success suddenly requires reaches into a reward zone that is centered 30° from the target. Alternatively, the reward zone can be shifted in a gradual manner, for example in 5° increments eventually reaching a maximum displacement of 30°. Following the introduction of the perturbation, success requires a movement that is off-target. While participants can find it challenging to learn when the shift is large or introduced abruptly (Brudner et al., 2016; Holland et al., 2018), many studies have shown that binary feedback is sufficient to produce learning when the shift is introduced in a gradual manner (Cashaback et al., 2019; Izawa & Shadmehr, 2011; Therrien et al., 2016, 2018; van der Kooij et al., 2021; van der Kooij & Smeets, 2018).

While sensory prediction errors and binary reward feedback can produce similar adjustments in behavior, there are marked differences in the representational changes associated with these two forms of learning (Morehead & Orban de Xivry, 2021; Therrien & Wong, 2022). For example, adaptation from sensory prediction errors is greatly attenuated when any delay is introduced between the movement and feedback, whereas adaptation from binary reward feedback is minimally impacted by delays up to a few seconds (Brudner et al., 2016; Schween & Hegele, 2017). In addition, the acquired behavior is more persistent following reward-based feedback compared to error-based feedback (Bao & Lei, 2022; Galea et al., 2015; Shmuelof et al., 2012; Therrien et al., 2016).

Learning processes can also be evaluated in terms of the degree to which they result in implicit and explicit changes in behaviour. A large body of literature has shown that adaptation from sensory prediction errors occurs in an automatic and implicit manner (Kim et al., 2018; Mazzoni & Krakauer, 2006; Morehead et al., 2017). Adaptation can also result from re-aiming, which is explicit and under volitional control. To date, less is known about implicit changes in behaviour in response to binary feedback. Following the convention in the adaptation literature, a strong probe of implicit learning is to focus on behavioral changes that persist when the feedback is eliminated and participants are reminded to reach directly to the target (Maresch, Mudrik, et al., 2021; Maresch, Werner, et al., 2021). When probed in this manner following reward feedback, a small aftereffect is observed. For example, following a shift of the reward zone of 25°, the average heading angle at the start of the aftereffect phase was around 5° (Holland et al., 2018). This suggests that reward-based learning is largely the result of a volitional change in strategy. Consistent with this hypothesis, disrupting explicit processes by introducing a secondary task attenuates learning from binary feedback (Codol et al., 2018; Holland et al., 2018). Nonetheless, the fact that there is an aftereffect, even if small, indicates binary feedback can induce implicit learning (Codol et al., 2018; Holland et al., 2018, 2019).

What might be the source of this implicit component? We can consider two, non-mutually exclusive hypotheses. The first hypothesis centers on the idea that the behavioral change resulting from binary feedback includes a contribution from implicit, use-dependent learning. As implied by the name, use-dependent learning refers to a movement bias towards frequently repeated movements (Diedrichsen et al., 2010; Huang et al., 2011; Marinovic et al., 2017; Mawase et al., 2017; Tsay et al., 2022; Verstynen & Sabes, 2011). Tracking the reward zone will result in movements that are shifted in a consistent direction relative to the visual target. In an aftereffect phase, a use-dependent bias would produce a residual implicit bias in this direction. Interestingly, the 3-4° aftereffect following training with binary feedback is similar in magnitude to that observed in studies of use-dependent learning that exclude errors in action selection (Tsay et al., 2022).

A second hypothesis is that binary feedback induces implicit recalibration of a sensorimotor map. Mechanistically, implicit recalibration could occur because the binary feedback alters the contingency between action plans and their associated movements. Feedback that indicates task success would strengthen the association between the goal to reach to a visual target and movements linked to that target, even if these are towards a reward zone that is displaced relative to the visual target. Feedback that indicates task failure would weaken this association. Compared to error-based learning, recalibration from reward feedback would appear to be much more limited given that the aftereffect following binary feedback is much smaller than that following cursor feedback for similar perturbation sizes (Bond & Taylor, 2015; Codol et al., 2018; Holland et al., 2018; Leow et al., 2018; Taylor & Ivry, 2014).

Here, we report the results of an experiment designed to assess these use-dependent learning and implicit recalibration hypotheses. Providing binary feedback only, we examined how participants learned to respond to either a small (7.5°) or large perturbation (25°) of the reward zone. For both groups, the perturbation was introduced in a gradual manner. Assuming that participants in the Small perturbation condition will have little awareness of the perturbation, this condition provides a strong test of the role of implicit processes in reward-based learning. In contrast, we assumed that participants in the Large condition would eventually adopt a strategy.

To assess implicit learning in both conditions, we measured reaching in an aftereffect phase in which all feedback was eliminated and participants were instructed to reach directly to the target. The implicit recalibration and use-dependent hypothesis both predict aftereffects in the Small and Large conditions. To compare the two hypotheses, we included two probe targets in the aftereffect phase, displaced by 15° from the training target location (Fig 1c). The inclusion of the probe targets allowed us to ask how implicit learning, if observed, generalized. By the implicit recalibration hypothesis, we would expect that reaches to the probe targets would be biased to a similar extent and in the same direction as reaches to the trained target. By the use-dependent hypothesis, we should observe that reaches to the probe targets would be attracted towards the trained movement. For the Small perturbation condition, the biases to the two probe targets should be in the opposite direction since the trained movement falls between the two probe locations. The predictions are less clear for the Large perturbation condition and will depend on the magnitude of learning. Biases to the two probe targets will be in the same direction if participants fully track the 25° shift of the reward zone. However, if the trained movement falls short of the reward zone, the biases will become less symmetric and even have opposite signs once the trained movement is less than 15°.

**Fig. 1.**
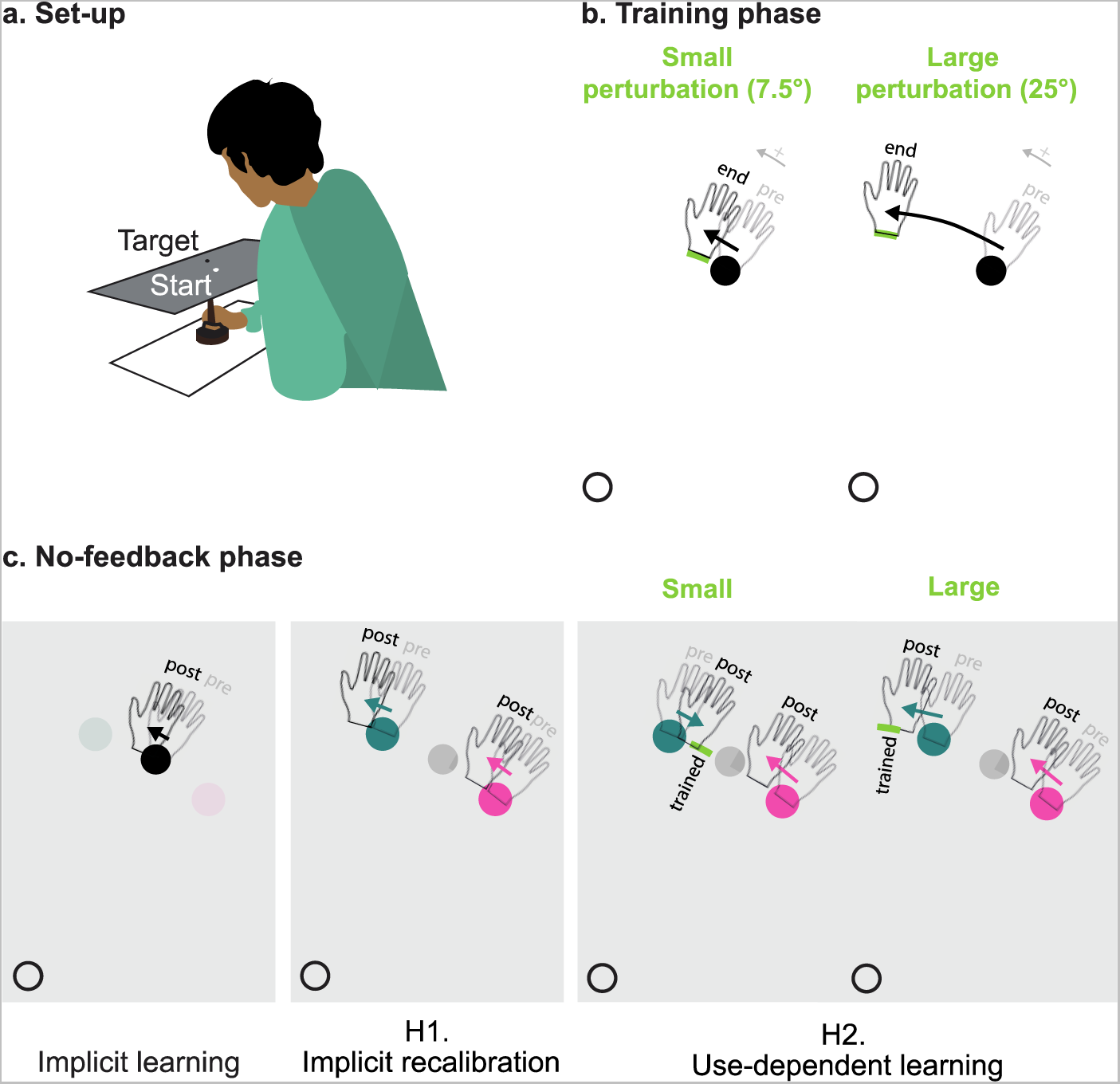
Schematic outline of key hypotheses on implicit reward-based motor learning. a. Schematic of a participant in the experimental apparatus. b. Training phase. Participants made center-out reaching movements from a white starting circle to a black training target. A pleasant auditory “ding” was provided when the movement passed within the reward zone (green arch); otherwise, an unpleasant “buzz” was played. c. No-feedback phase. Participants were instructed to reach directly to a visual target. The target appeared at the training location or one of two probe locations (+-15°). Participants were instructed to forgo any strategy adopted during the training phase. Left panel shows implicit learning as measured by an aftereffect, defined as a change in hand angle for reaches to the training target from pre-training (translucent hand) to post-training (solid hand). Middle panel shows probe target reaching predictions for the implicit recalibration hypothesis. Reaches will be biased in same direction for both probe targets independent of size of the perturbation. Right panel shows probe target reaching predictions for the use-dependent learning hypothesis. For the Small perturbation condition, the biases will be in opposite directions since the reaches during training fall between the two probe locations. For the Large perturbation condition, the direction of the bias for the probe target nearest the reward zone will depend on the degree of learning (example here is for a participant who shows full learning)

## Methods

### Participants

68 right-handed young adults were recruited from the research participant pool of the Department of Psychology at the University of California, Berkeley. 28 (22 females, 6 males; reported age: mean 20.5, SD 2.3 years) were assigned to the “Small” perturbation group and 40 (27 females, 13 males; reported age: mean 21.5, SD 5.7 years) were assigned to the “Large” perturbation group. Participants received either course credit or financial compensation for their participation, along with a $5 completion bonus paid to all participants. Based on self-reports, participants had normal or corrected-to-normal vision and hearing. The protocol was approved by the institutional review board at UC Berkeley.

Of the original 68 participants, 20 were excluded from all analyses. 16 of these (8 per group) were excluded based on their responses to a post-experiment questionnaire (see “Experimental design”) that indicated they failed to follow the instructions. Four other participants in the Large group were excluded for idiosyncratic reasons: One fell asleep during the task, one reported, after the experiment having performed in a similar experiment, one did not use the apparatus correctly, and one experienced an equipment failure. Thus, the analyses reported below are based on data obtained from 20 participants (16 females; 10 for credit; reported age: mean 20.9, SD 2.4 years) in the Small perturbation group and of 28 participants (16 females; 16 for credit; reported age: mean 21.8, SD 6.0 years) in the Large perturbation group.

### Experimental set up

The participant sat in front of a table in a small, darkened room. A horizontally-oriented computer screen (24”, ASUS, Taipei, Taiwan) constituted the upper surface of the table, with a 17” digitizing tablet (Wacom Co., Kazo, Japan) positioned 27 cm below the screen (Fig 1a). Stimuli were presented on the computer (refresh rate = 60 Hz) and the participant’s movements along the digitizing tablet were recorded from a digitizing pen (sampling rate = 200 Hz) that was embedded in a custom-made paddle, ensuring the pen maintained a vertical position. Vision of the hand was obscured by the screen. A computer (Dell OptiPlex 7040, Round Rock, Texas) with a Windows 7 operating system (Microsoft Co., Redmond, Washington) was used to run the custom experimental software in Matlab (The MathWorks, Natick, Massachusetts), using Psychtoolbox extensions (Brainard, 1997; Kleiner et al., 2007).

### Trial structure

Each trial started with the appearance of a white “start” circle (radius = 0.42 cm), presented near the center of the screen. The participant was required to move the paddle to position the digitizing pen within the start circle. To guide the participant to the start location, a white ring was presented, with the radius of the ring indicating the distance from the pen to the start position. Movement towards the start position reduced the size of the ring. When the pen was within 0.84 cm of the start circle, the ring was replaced by a white circle (radius = 0.17 cm) that indicated the position of the pen, allowing the participant to move the pen into the start circle.

When the paddle had been in the start circle for 300 ms, a visual target (circle with radius = 0.28 cm) appeared 7 cm from the start circle at either 45°, 60° or 75° (Figure 1b, c). The participant was instructed to move in rapid manner, attempting to slice through the target. Auditory feedback was presented when the movement amplitude exceeded 7 cm. On trials with performance feedback (see below), a pleasant “bing” indicated that the movement was successful (e.g., passed through the target when feedback was veridical) and an aversive “buzz” indicated that the movement was unsuccessful. On no feedback trials (in the baseline and aftereffect phases), a “knock” sound was played. This indicated that the required reach amplitude had been exceeded but it did not provide feedback on whether the movement was within the reward zone or not. To make participants move at similar, and relatively rapid speeds, 800 ms after the performance feedback an auditory message “Too slow” was played if the movement time was longer than 600 ms. This was the case on 3% of the trials. We did not exclude these slow trials from the analyses.

The feedback ring appeared directly after the feedback was given. Note that by using a ring during the return movement, the participant received feedback indicating only the radial position of the hand. Angular position was only provided when the hand was very close to the start position: then, the ring turned into a cursor. This method was used so that any effect of adaptation to the rotated feedback (see below) would be minimally visible to the participant during the return movement.

### Experimental design

The experimenter instructed the participant that the purpose of the experiment was to study how well people can control arm movements in the absence of visual feedback. The participant was told that they would control an invisible cursor, and they were asked to make reaching movements that would make the invisible cursor intersect a visual target (Fig 1a). The experimenter described how a “bing” and “buzz” would indicate if the reach had intersected or missed the target, respectively. The experimenter then completed 10 demonstration trials to demonstrate how the hand controlled the cursor movement. The target was always presented at the 60° location and during these trials, the auditory feedback was accompanied by veridical cursor feedback.

After the ten demonstration trials, the participant was told that the cursor would no longer be visible during the reach, but that auditory feedback would be presented on most trials to indicate task outcome. However, on some trials, the participant would hear a “knock” sound, and this sound was uninformative concerning task outcome. To motivate the participant for all trials, the participant was informed that the computer kept track of all successful reaches and that a score in the top-third of high scores across participants would result in a $5 bonus (which was actually paid to all participants).

The main experiment consisted of three phases: baseline, training, and aftereffect, with the experimenter providing instructions at the beginning of each phase. The baseline phase was composed of 150 trials with feedback limited to the uninformative “knock” sound. The target appeared at each of the three possible locations on 50 of the baseline trials, with the order determined randomly. These trials allowed the participant to become familiar with the apparatus, learn to move at the appropriate speed, and provided a measure of natural biases for each of the three target locations (Kuling et al., 2019; van der Kooij et al., 2013).

The training phase was composed of 700 trials, with the target always appearing at the middle location (60°) and auditory feedback provided to indicate target hits or misses. For the first 100 trials, the reward zone was centered around the participant’s individual bias while reaching to the trained target and extended 2° in both directions; if, for example, the individual’s mean reach to the central target was rotated by 3° in the clockwise direction (at 57°), the initial reward zone spanned from 55° – 59°. Unbeknownst to the participant, the reward zone was gradually shifted over the next 500 trials. This was achieved by rotating the reward zone by 1.5° every 100 trials for the Small perturbation group and by 2.5° every 50 trials for the Large perturbation group. The rotation was either clockwise or counterclockwise, counterbalanced between participants. For the last 100 trials of the training phase, the reward zone remained fixed, displaced by 7.5° or 25° from its starting position for the Small and Large perturbation groups, respectively. A two-minute break was provided halfway through the 700-trial training phase.

Note that we expected that the participants in the Small group would likely remain unaware of the perturbation since the shift was introduced gradually and the total displacement fell within 1-2 standard deviations of normal reach variability (Gaffin-Cahn et al., 2019). In contrast, we expected that participants in the Large group would likely become aware of the perturbation at some point during the training phase as the discrepancy between the visual target and hand movement would likely fall outside the individuals’ normal reach variability.

Following the training phase, the participant completed an aftereffect phase of 150 trials. Prior to the start of the phase, the participant was instructed that the feedback might have been altered over the course of the training phase. To equally inform and instruct participants with different levels of awareness of the perturbation, the participant was informed that there were two groups of participants, an aligned group and a misaligned group. For the aligned group, the invisible cursor had always moved exactly with the position of the hand; for the misaligned group, the invisible cursor was slightly displaced from the position of the hand. To ensure that the participant understood the difference, they were asked to explain the difference between the two groups in their own words. If the explanation failed to capture the difference, the experimenter repeated the explanation. The experimenter then stated that for the final phase of the experiment, the cursor would be aligned with the hand for everyone, irrespective of initial group assignment and thus, they should reach straight to the target to make the cursor hit the target. As in the baseline phase, reaches during this phase were performed with only the uninformative feedback, with the phase composed of 50 reaches to each of the three targets. Participants were again instructed that accuracy would be recorded during this phase to determine a final performance bonus.

At the end of the experiment, the participant completed a questionnaire consisting of five questions (Online resource 1). Question 1 asked if they believed the feedback had been veridical or perturbed and Question 2 asked for their confidence concerning their response to Question 1, using a 7-point rating scale (1= no confidence, 7= very confident). For Questions 3 and 4, the participants were asked to report (forced choice) where they had aimed during the training and aftereffect phases, respectively. Note that Question 4 was used to determine if the participant had followed the instructions. Those who answered that they had aimed to the left or right of the target during the aftereffect phase were excluded from all of the analyses (n=16). For Question 5, the participant was informed that they had been in the Misaligned feedback group and were asked to indicate (forced choice) if the feedback had been perturbed: to the left or to the right. As the answers to this question were below chance level in the Small perturbation group, for the Large perturbation group, the illustrations for the two choices were slightly changed (Online resource 1) to match the hand movements of the participants better.

The total duration of the experiment was approximately one hour.

### Data analysis

Reach angle was determined by the line from the start position to where the digitizing pen crossed the 7 cm radius around the start position. The mean reach angle during the baseline trials was used to characterize individual biases for each of the three target locations separately (50 reaches/target). All analyses were based on the reach angles during the training and aftereffect phases, with these angles expressed relative to that participant’s baseline bias for the corresponding target. Positive values correspond to reach angles shifted in the direction of the rotated reward zone.

We calculated final learning as the mean reach angle in the last 100 trials of the training phase. To test for implicit learning, we calculated the mean reach angle to the training target in the aftereffect phase. For generalization, we calculated the mean reach angle for each of the two probe targets in the aftereffect phase.

#### Statistics

A preliminary analysis indicated that the final learning and aftereffect scores were not normally distributed (see Fig 2). Therefore, we employed non-parametric tests in the statistical evaluation of the results. To test whether the final learning and aftereffect were larger than zero, we performed a one-tailed Wilcoxon signed-rank test on these variables for each group (Small and Large). To test whether implicit learning was different for the two perturbation sizes, we performed a two-tailed Wilcoxon rank-sum test on the aftereffect scores for the two groups.

**Fig. 2.**
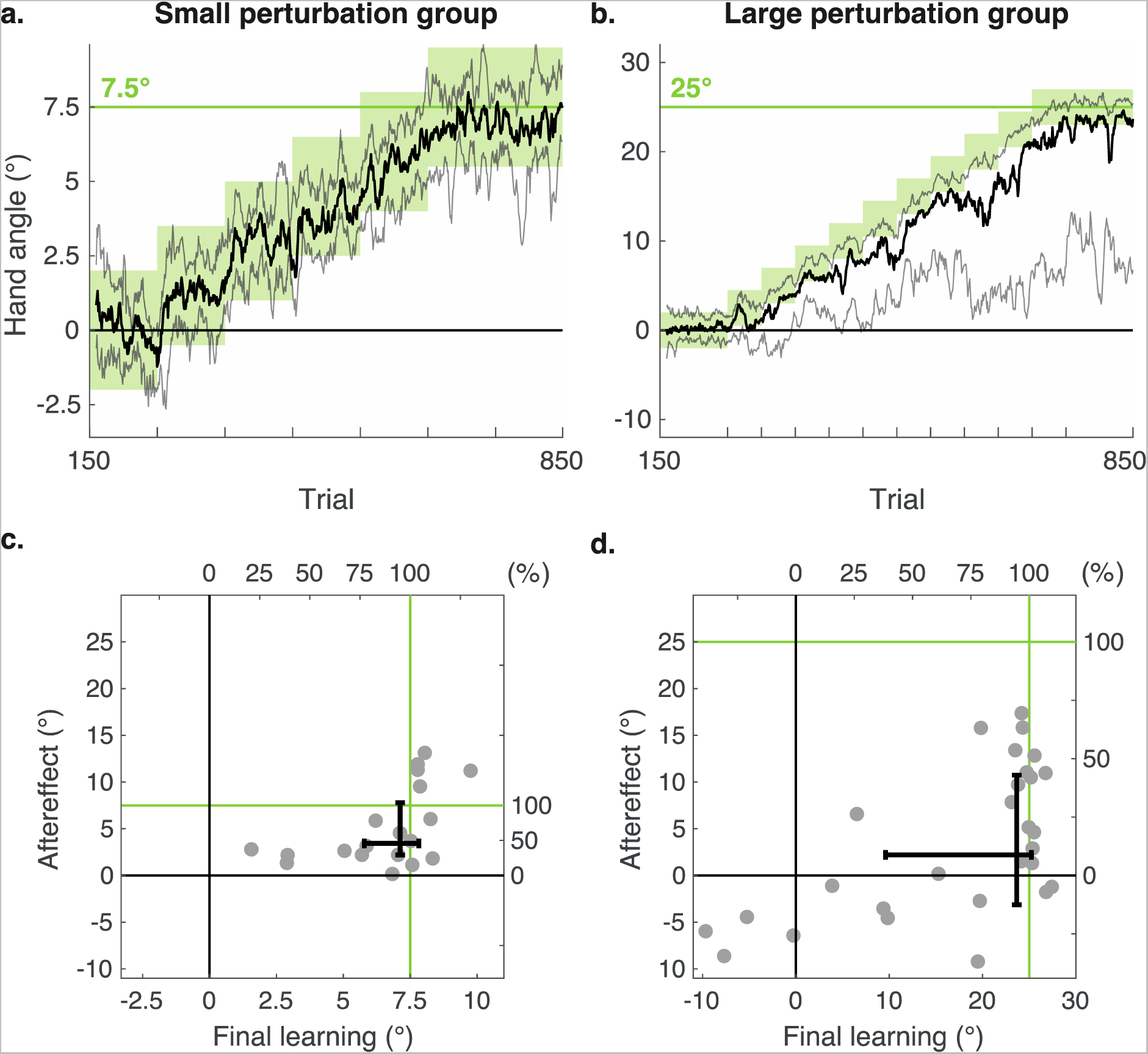
The effect of binary reward feedback on reaching. a, b. The training phase. Gradually changing the rewarded hand angles (green zone) leads to learning, as indicated by the change in reach angle. We plot the median (solid thick lines) over all participants with the interquartile range (opaque lines) for the Small perturbation group (a) and Large perturbation group (b). Note that the vertical axes are scaled to the perturbation size. For display purposes, the curves are smoothed with a running average with a window size of 10 trials. c, d. Aftereffect as a function of the final learning for both groups. Each grey dot corresponds to a participant and error bars indicate the interquartile range per group. The green lines indicate perturbation size

For the generalization data, we defined the percentage generalization as the mean of the two probe target biases, divided by the aftereffect at the training target. We used a one-tailed Wilcoxon signed-rank test to test whether the percentage generalization values were significantly larger than zero. To evaluate the form of generalization, we defined generalization asymmetry as the difference between the reaching bias to the probe target opposite the reward zone and the probe target in the direction of the reward zone. The use-dependent learning hypothesis predicts that this value will be positive for the Small perturbation condition. The implicit recalibration hypothesis predicts that this value will be zero (if generalization is exactly the same for both targets, but see (Nikooyan & Ahmed, 2015)). To evaluate the two hypotheses, we used a Wilcoxon signed-rank test to test whether the generalization asymmetry values were significantly greater than zero.

No statistics were performed on the questionnaire data.

## Results

### Learning

To evaluate how people modified their movements given the gradual change in the reward zone, we analyzed the reach angle at the end of learning in both the Small (max shift of 7.5°) and Large (max shift of 25°) perturbation groups. Both groups learned to compensate for the feedback perturbation (Fig 2a,b). Participants in the Small perturbation group showed a median final learning of 7.1° (IQR [5.8°, 7.8°], p < 0.01, z = 3.9, Ws = 210, r = 0.20) (Fig 2c, horizontal axis). Participants in the Large perturbation group showed a median final learning of 23.7° (IQR [9.6°, 25.2°], p < 0.01, z = 4.2, Ws = 390, r = 0.16 (Fig 2d, horizontal axis). For both groups, this corresponds to a mean percentual change of 95% of the perturbation size (Small: IQR = 77% −104%; Large: IQR = 38% −101%).

As can be seen in Fig 2c,d (horizontal axes), learning was more variable in the Large perturbation group than in the Small perturbation group. For the latter, all of the participants changed their reaches in the direction of the perturbation and 86% ended up with a mean heading angle over the final 100 trials that was within the final reward zone. In contrast, only 70% of the participants in the Large perturbation group reached the final reward zone (Online resource 3). Four participants in this group exhibited a mean hand angle over the final 100 trials that was in the opposite direction of the reward zone.

### Aftereffect

The central aim of our study was to examine whether binary feedback regarding success or failure induces implicit motor learning. To this end, we focused on the reach direction during the aftereffect phase when the feedback was removed and participants were instructed to reach directly to the target.

Both groups showed a significant aftereffect (Fig 2c, d vertical axes). Participants in the Small perturbation group had a median aftereffect of 3.4° (IQR [2.2°, 7.8°]; p < 0.01, z=3.90, Ws = 210, r = 0.20). Participants in the Large perturbation group had a median aftereffect of 2.2° (IQR [-3.1°, 10.7°], p < 0.05, z = 2.02, Ws = 292, r = 0.07). Importantly, we found no difference between the magnitude of the aftereffect for the Small and Large perturbation groups (p = 0.24, z=-1.2, U = 434).

In summary, the aftereffect data indicate that there is an implicit component to learning that occurs in response to binary feedback. The magnitude of the aftereffect in both the Small and Large perturbation groups was of similar size and quite small.

### Generalization

We included reaches to two probe targets in the aftereffect phase, asking how learning generalized to regions of the workspace neighboring the training target. Both groups exhibited generalization in that the reaches to the probe locations were significantly shifted from the baseline phase. In terms of the direction of the shift, the mean values were all positive, meaning that the change in reach direction for the probe targets was in the same direction as the change in reach direction to the training target (Fig 3a). Participants in the Small perturbation group had a median reaching bias of 3.5° to the probe target in the direction of the learning and of 3.6° to the other probe target (Online resource 4). The corresponding biases were 1.6° and 0.7° for the Large perturbation group.

**Fig. 3.**
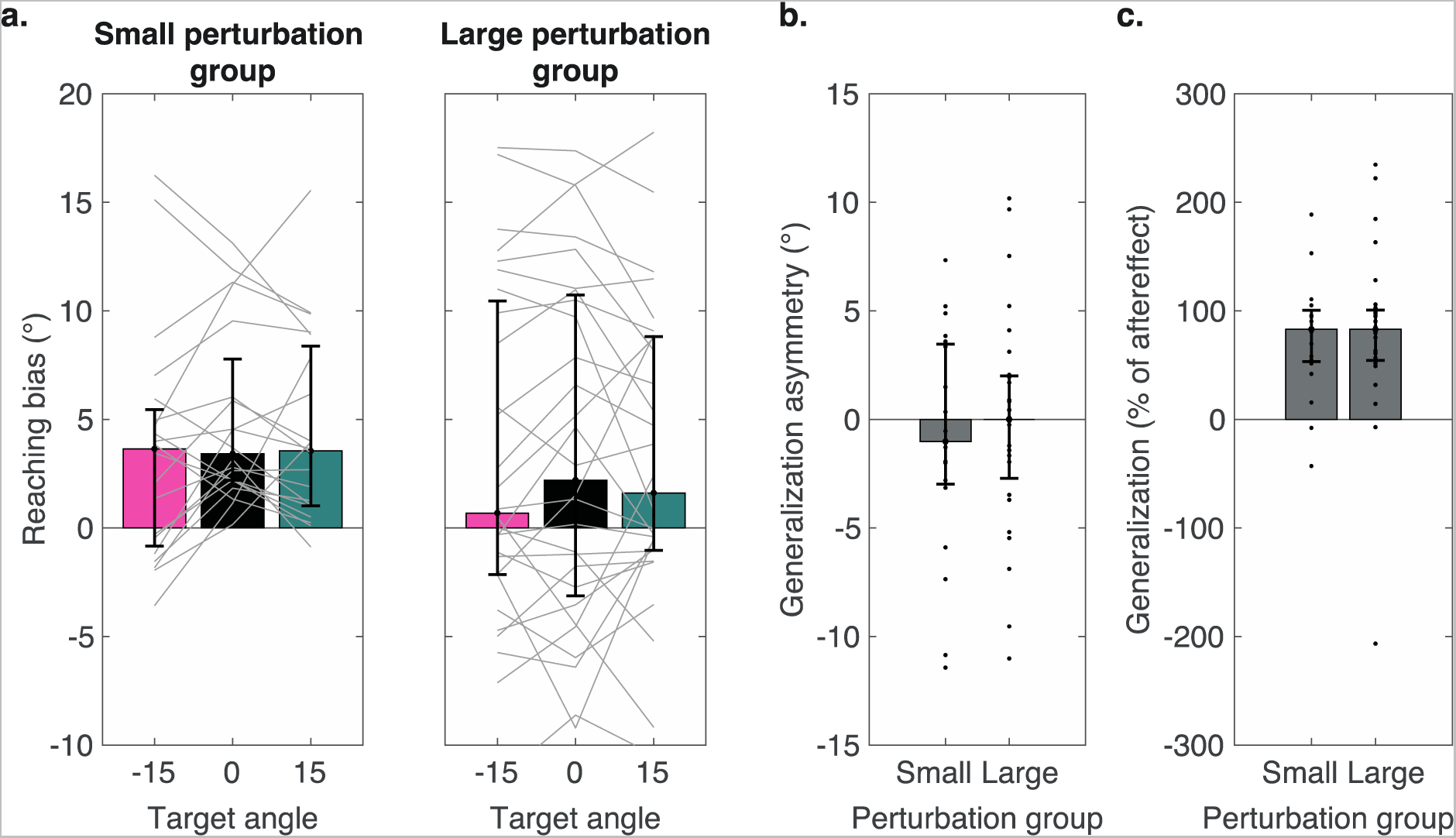
Aftereffects and generalization of learning. Bars and error bars indicate medians and interquartile ranges. a. Reaching biases for the training target (black) and two probe targets (see Fig 1c). Thin lines indicate data from individual participants. b. Asymmetry in reaching biases to probe targets. Dots indicate the individual participants in the two groups. c. Generalization quantified as a percentage of the aftereffect. The participants with large positive and negative values are the ones with a small aftereffect

The generalization data are consistent with the implicit recalibration hypothesis. For each participant, generalization (Fig 3c) was calculated as the mean of the two probe target biases as a percentage of the aftereffect. We found significant generalization of 83% of the aftereffect in both the Small (IQR [53.3%, 100.5%], p < 0.01, z = 3.7, Ws = 205, r = 0.19) and Large (IQR [54.3%,100.6%], p < 0.01, z = 4.0, Ws = 379, r = 0.14) perturbation group.

The generalization data are not consistent with the use-dependent learning hypothesis. The use-dependent learning hypothesis had predicted biases in opposite directions for the two probes in the Small perturbation group since the trained movement was between the two probe targets. This would predict positive generalization asymmetry scores. In the Large perturbation group, predictions were less clear since they depend on the location of the trained movement relative to the probe targets. For participants who fully followed the reward zone, the trained movement was beyond both probe targets; for others, the trained movement was either between or beyond both probe targets. For both groups, the analyses showed that the asymmetry scores were not significantly larger than zero (Small: median = − 1.0°, IQR [-3.0°, 3.5°], p = 0.55, z = −0.06, Ws=89; Large: median = 0.0°, IQR [-2.7°, 2.0°], IQR [53.3%, 100.5%], p = 0.96, z = −0.05, Ws=201). These null results, coupled with the fact that we did observe significant generalization, are consistent with the implicit recalibration hypothesis.

### Awareness of the feedback perturbation

As expected, participants in the Small perturbation group were generally unaware that the reward zone had shifted over the course of the experiment. When asked to judge if they had been in the group with veridical feedback or shifted feedback, 60% reported that the feedback was not perturbed with an average confidence of 3.3 on a 7-point scale (Online resource 1, 3). When forced to choose between saying if they aimed left, right, or straight to the target during the training phase, 50% reported having aimed straight to the target and 50% reported aiming away from the target. However, of the latter, half reported aiming in the direction of the shifted reward zone and the other half reported aiming in the opposite direction. These survey data, in combination with the fact that all participants in the Small perturbation group showed a shift in reaching in the direction of the perturbation, provide compelling evidence that there was little if any awareness of the experimental manipulation nor use of a compensatory strategy.

A very different picture emerged from the survey data for the Large perturbation group. The majority (82%) reported that the feedback was perturbed with an average confidence of 4.8 on the 7-point scale. When asked whether they aimed left of, right of or straight to the target during the training phase, 75% of the participants reported having aimed off target in the direction of the shifted reward zone, whereas 21% reported having aimed straight to the target. In summary, the survey data indicate that the participants in the Large perturbation group were aware of the experimental manipulation and adopted a re-aiming strategy to compensate for the shift in the reward zone.

There was no clear relation between the questionnaire reports and aftereffects (Online resource 2).

## Discussion

In the present study, we examined whether binary feedback can induce implicit learning in response to shifts in a hidden reward zone. Based on previous work (Codol et al., 2018; Holland et al., 2018, 2019), we expected that the learning would include an implicit component. Participants performed a center-out reaching task and were only provided binary feedback to indicate if the movement ended in a reward zone that gradually shifted to be centered 7.5° or 25° from the visual target, with the expectation that awareness of the perturbation would be minimal in the former and that the latter would entail some explicit component. During training, participants in both groups learned to compensate for the rotated feedback. When the feedback was removed after training and participants were instructed to move to the target, their reaches were biased in the direction of learning, with an aftereffect of 2-3° in both groups. To test generalization, the no-feedback phase also included reaches to probe targets that flanked the training target. On these probe target trials, participants exhibited a shift in reach angle that was in the same direction as the shift associated with the training target. These results suggest that binary feedback can induce implicit reward-based motor learning and that this learning reflects implicit recalibration of a sensorimotor map.

### Small and saturated implicit learning in response to binary feedback

Our study employed multiple approaches to prevent explicit processes from contaminating our assessment of implicit learning. First, we focused on the aftereffect in a phase without feedback and in which we provided explicit instructions to stop using any strategy that might have been used during the training. Second, we introduced the perturbation in a gradual manner, and most importantly, included a small perturbation group in which the displacement per step was within 1.5 standard deviations of baseline reach variability (Online resource 4) (Gaffin-Cahn et al., 2019). Thus, for this group, it is likely that behavioral changes during the training phase occurred implicitly. Third, we used questionnaires to directly assess awareness of the perturbation. The responses to the survey confirmed that, during the perturbation phase, awareness and strategy use were minimal in the Small perturbation group but high in the Large perturbation group.

We observed a small, but consistent aftereffect of around 2-3° in both the Small and Large perturbation groups. The magnitude of this effect for the latter group is smaller than that previously reported in other studies using a perturbation of comparable size; for example, in Holland et al. (2018, see also, 2019), the aftereffects in response to a perturbation of 25° were around 5°. However, during their no-feedback aftereffect phase, Holland et al. first instructed participants to keep reaching as they had done during training. After this phase, they were instructed to stop using a strategy. This protocol may have contaminated the final aftereffect measure by adding extra strategy trials and the challenge to switch between tasks.

The inclusion of the Small perturbation group not only provided a condition in which awareness should be minimized during the training phase, but also allowed us to directly compare how perturbation size impacted the magnitude of implicit learning from binary feedback. Interestingly, the size of the aftereffect did not scale with perturbation size. Indeed, in terms of mean value, the size was larger in the Small condition (3.4°) compared to the Large condition (2.2°), although this difference was not significant. While future testing is required to sample a broader range of perturbation sizes, the present results suggest that the magnitude of implicit learning from binary feedback is relatively small and saturates, at least for perturbations larger than 7.5°.

### Mechanisms of implicit learning in response to binary feedback

In the following section, we will consider the mechanisms underlying implicit learning in response to binary feedback. Similar to what has been reported in studies of error-based learning (Bond & Taylor, 2015; Morehead et al., 2017) and use-dependent learning (Tsay et al., 2022), implicit learning in response to binary feedback seems to saturate. However, there are notable differences between these three implicit forms of learning. While the magnitude of implicit use-dependent biases is similar to the magnitude of the aftereffect observed in the present study, the generalization pattern did not show any evidence of attraction towards the training location. As such, the current results fail to support the idea that implicit learning from binary feedback is a manifestation of use-dependent learning. On the other hand, while the generalization pattern is similar for binary and cursor feedback, the magnitude of the binary feedback effect is much smaller than that observed in response to cursor feedback (Bond & Taylor, 2015; Morehead et al., 2017). This size discrepancy makes it unlikely that binary feedback operates on similar mechanisms in inducing implicit recalibration of the sensorimotor map.

How, then, does binary feedback result in implicit learning? We outline three implicit recalibration hypotheses. First, implicit learning in response to binary feedback could be the result of motor recalibration, retuning the mapping between a visual target location and its associated movement. The contingency between action and reward outcome will lead to that action being associated with a new movement plan (Avraham et al., 2022). This hypothesis predicts that there is no sensory recalibration: training would not influence reports of where the visual target is perceived and perceived locations of the hand so that they are similar before and after training. Second, implicit learning could be the result of visual recalibration of the target, i.e., a bias in the perceived location of the visual target. This hypothesis predicts visual sensory remapping: for example, if asked to report the perceived target location by reaching with the non-trained hand, we would observe a bias towards the reward zone (Simani et al., 2007). Third, implicit learning could be the result of proprioceptive recalibration, i.e., a bias in perceived hand position. This hypothesis predicts proprioceptive sensory remapping. For example, static reports of perceived hand position would be biased in the opposite direction of the perturbation (Tsay & Ivry, 2022).

Future studies employing fine-grained psychophysical tests can evaluate the merits of these different hypotheses, asking if implicit learning in response to binary feedback originates from implicit recalibration of a sensory and/or motor mapping, and how this evolves over the course of learning.

## Conclusion

Our data add to a growing body of evidence indicating that motor learning encompasses multiple processes where both explicit and implicit processes drive behavioral changes (Kim et al., 2021; Morehead & Orban de Xivry, 2021; Therrien & Wong, 2022). The results provide compelling evidence of implicit learning in response to binary feedback and rule out that this effect is a form of use-dependent learning. Less clear is whether this implicit learning entails the same mechanisms, albeit in attenuated form, as occur during learning from sensory prediction errors, or reflects the operation of distinct, reward-based mechanisms.

## Supporting information

Online resource 1. Questionnaires

Online resource 2. Questionnaire and behavioural results

Online resource 3. Learners and non-learners

Online resource 4. Reach angle variability

## Acknowledgments

The research was funded by the Nederlandse Organisatie voor Wetenschappelijk Onderzoek, Toegepaste en Technische Wetenschappen Open Technologie Programma (NWO-TTW OTP grant 15989), and by the United States National Institutes of Health (NIH grants R35NS116883 and NS105839). The funders had no role in study design, data collection and analysis, decision to publish, or preparation of the manuscript.

## Authorship

Conceptualization: Nina M. van Mastrigt, Rich Ivry, Tianhe Wang, Jonathan Tsay, Guy Avraham

Funding acquisition: Jeroen B. J. Smeets, Katinka van der Kooij

Investigation: Nina M. van Mastrigt.

Methodology: everyone

Software: Tianhe Wang (PsychToolbox), Nina van Mastrigt (Matlab + psychtoolbox)

Supervision: Rich Ivry, Jeroen B. J. Smeets, Katinka van der Kooij.

Visualization: Nina M. van Mastrigt

Writing – original draft: Nina M. van Mastrigt

Writing – review & editing: everyone

## Data and code availability

Data and code can be accessed on the Open Science Framework (https://osf.io/x7hp9/).

## Supplementary information

### Online resource 1. Post-experiment questionnaire

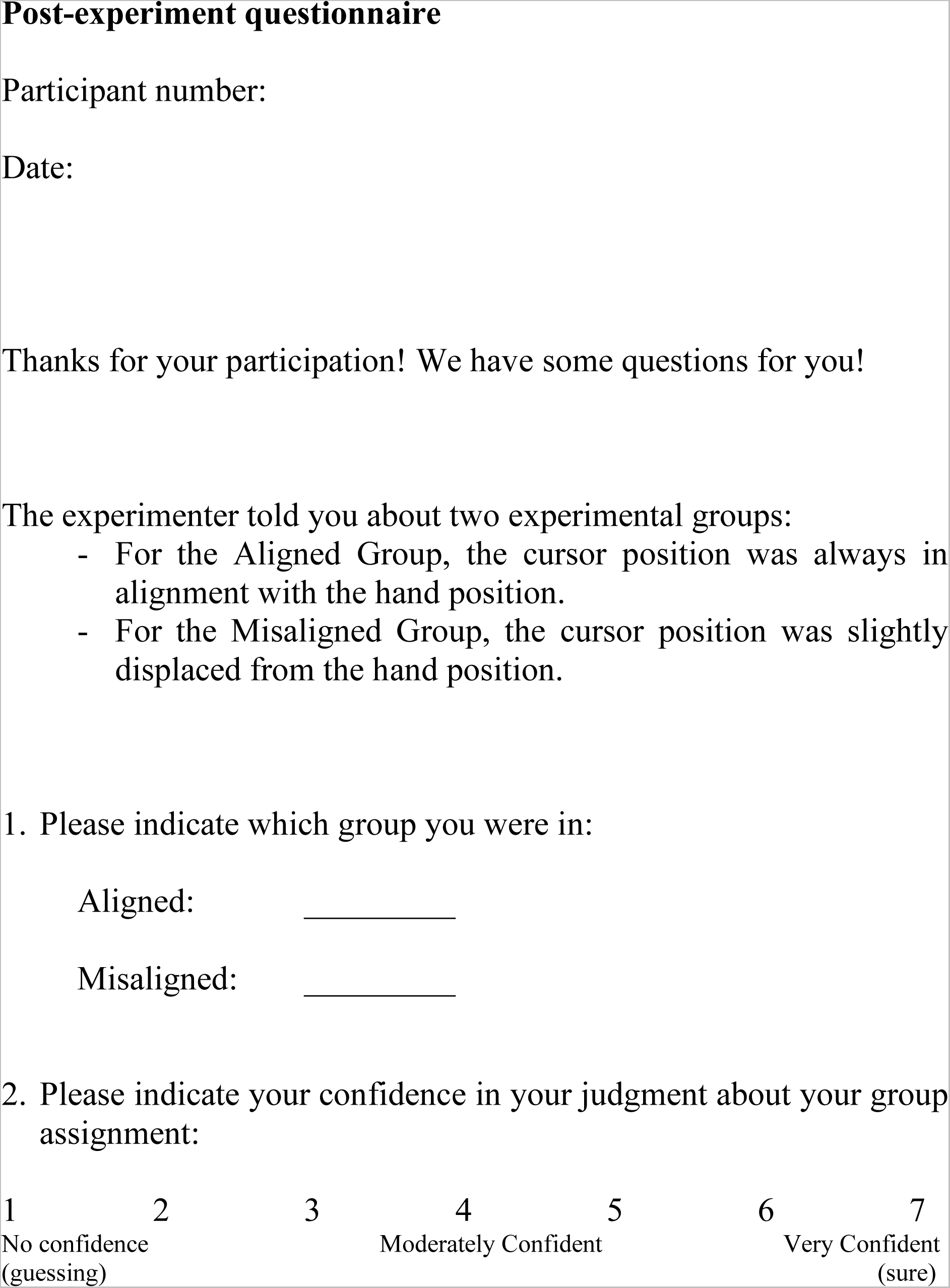
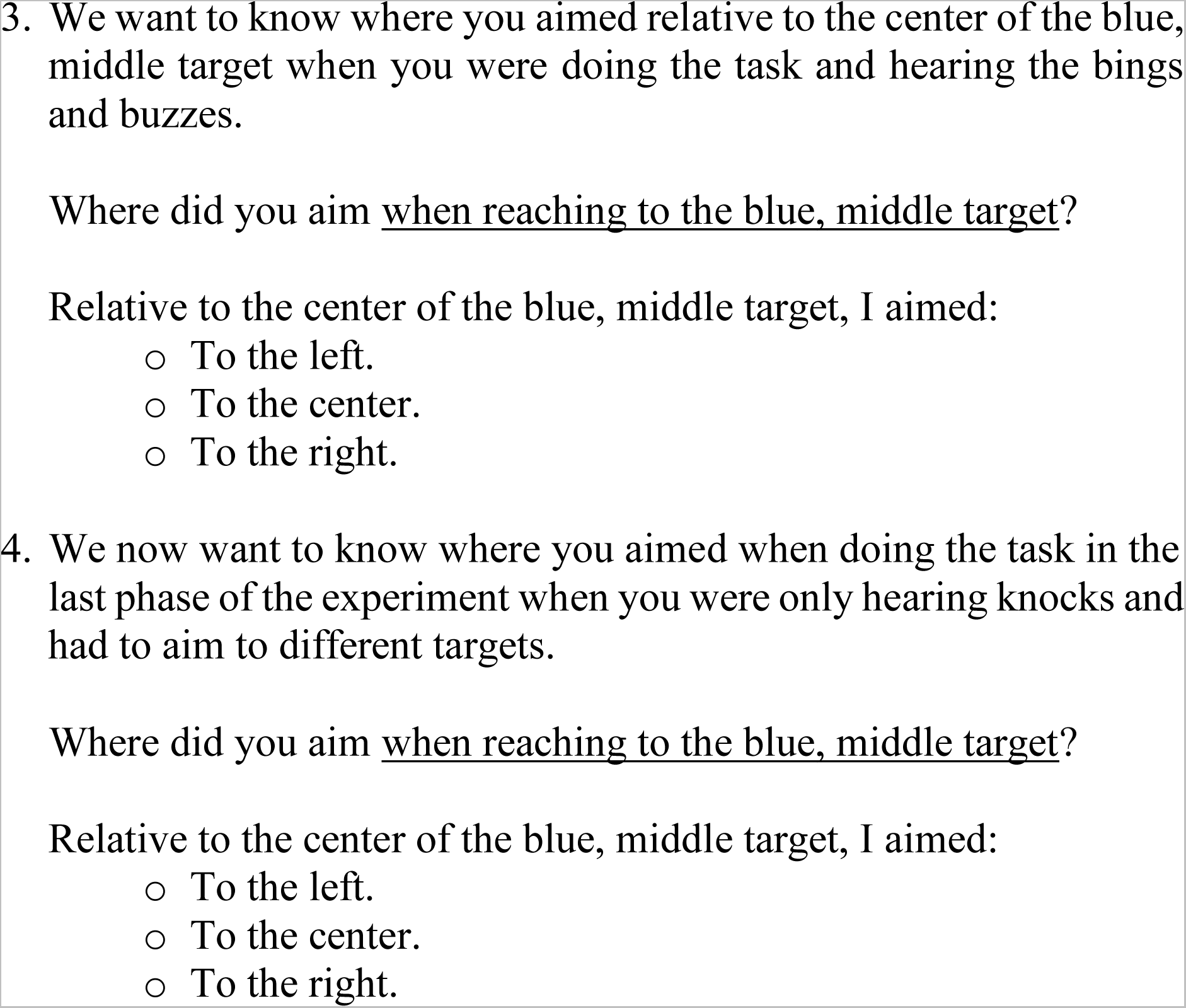
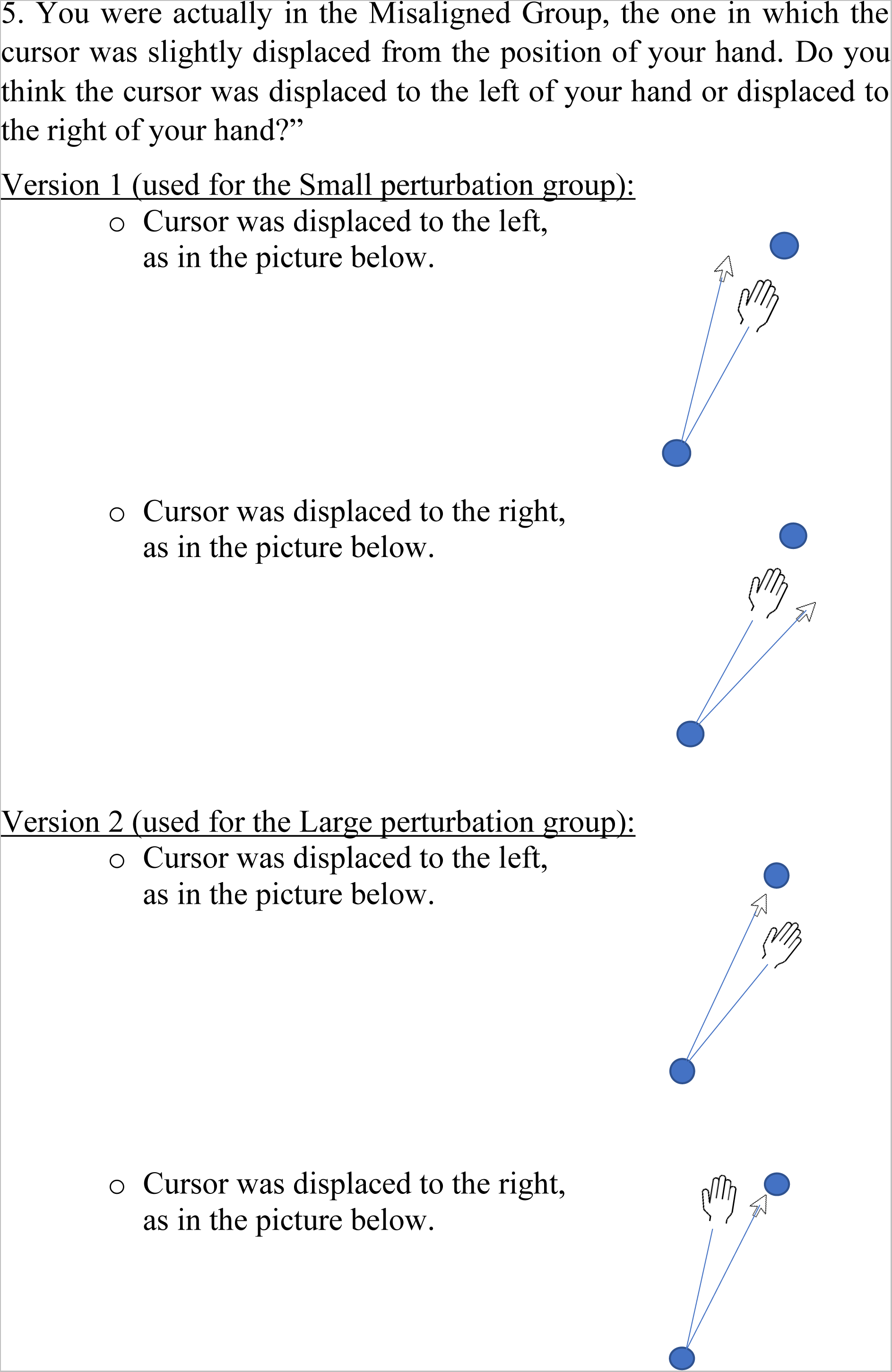

### Online resource 2. Questionnaire results

**Online resource 2.**
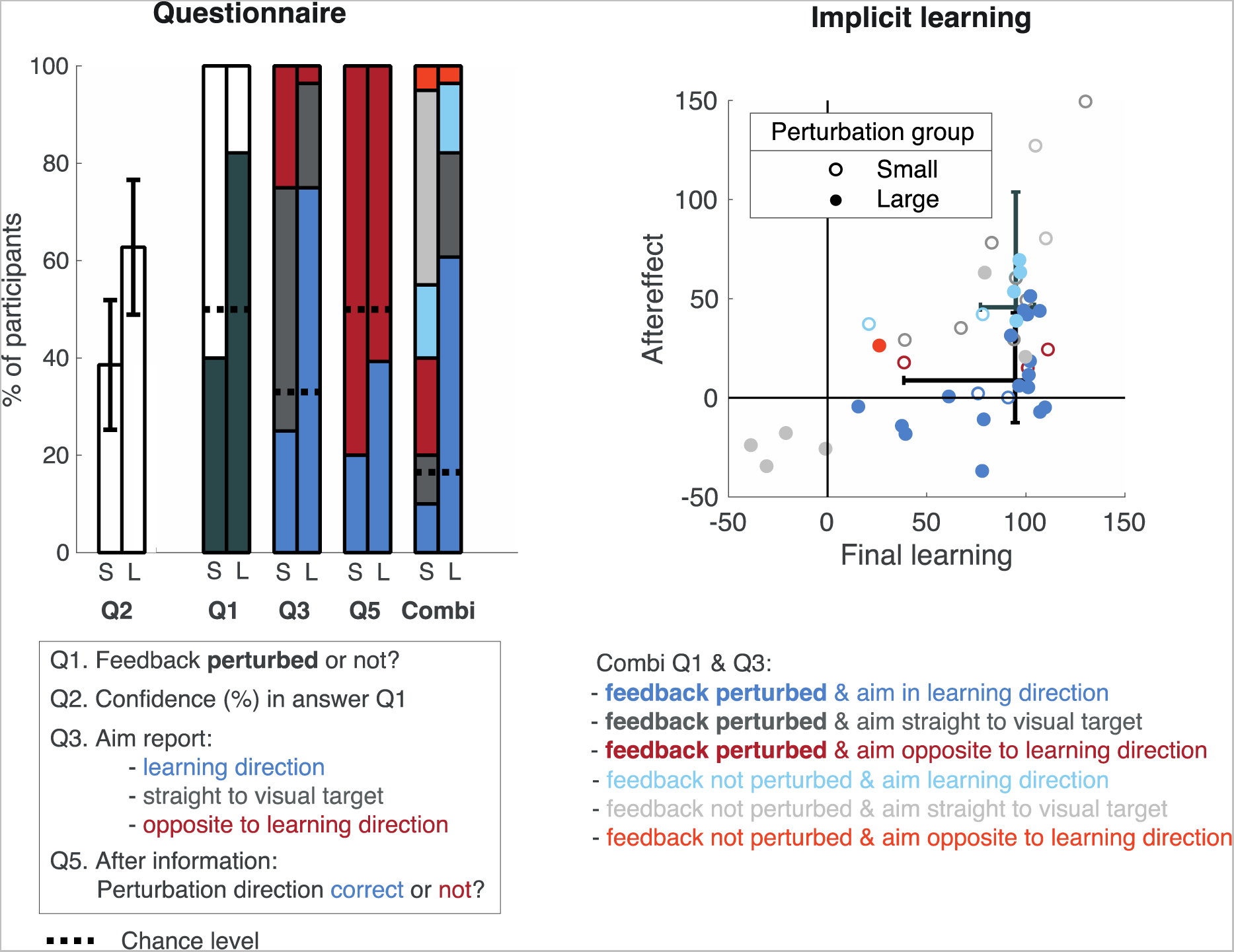
Questionnaire results related to learning for the Small Perturbation group (left bars) and the Large Perturbation group (right bars). Q’s correspond to question numbers on the questionnaire. See Online resource 1 for the post-experiment questionnaires for the Small perturbation group and Large perturbation group.

### Online resource 3. Learners and non-learners

**Online resource 3.**
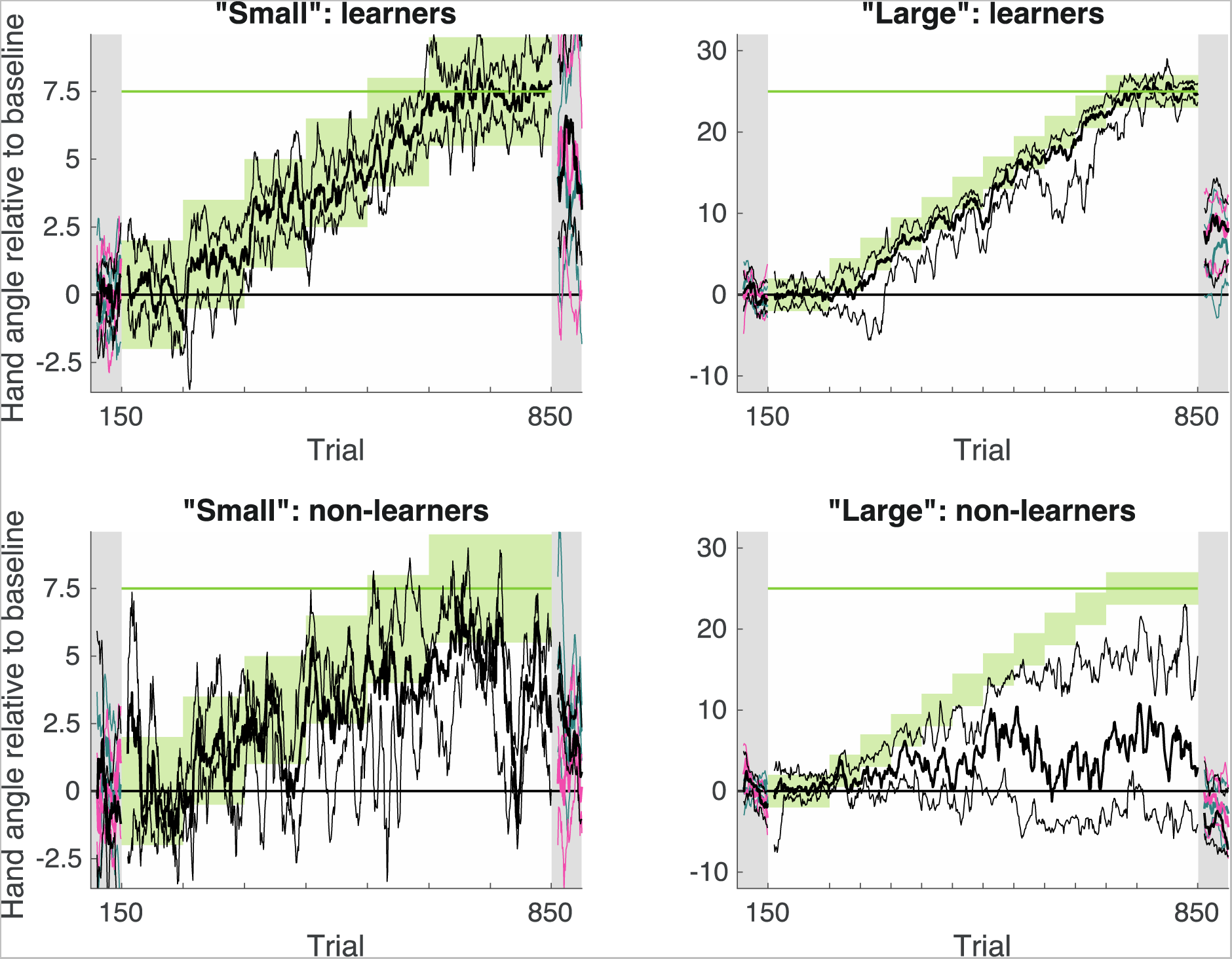
Learners and non-learners. Participants were divided into two groups based on whether final learning was in the reward zone (“learners”, top rows) or not (“non-learners”, bottom rows). By this definition, there were 16 learners and 4 non-learners in the Small Perturbation group (left) and 16 learners and 12 non-learners in the Large Perturbation group.

### Online resource 4. Reach angle variability

**Online resource 4.**
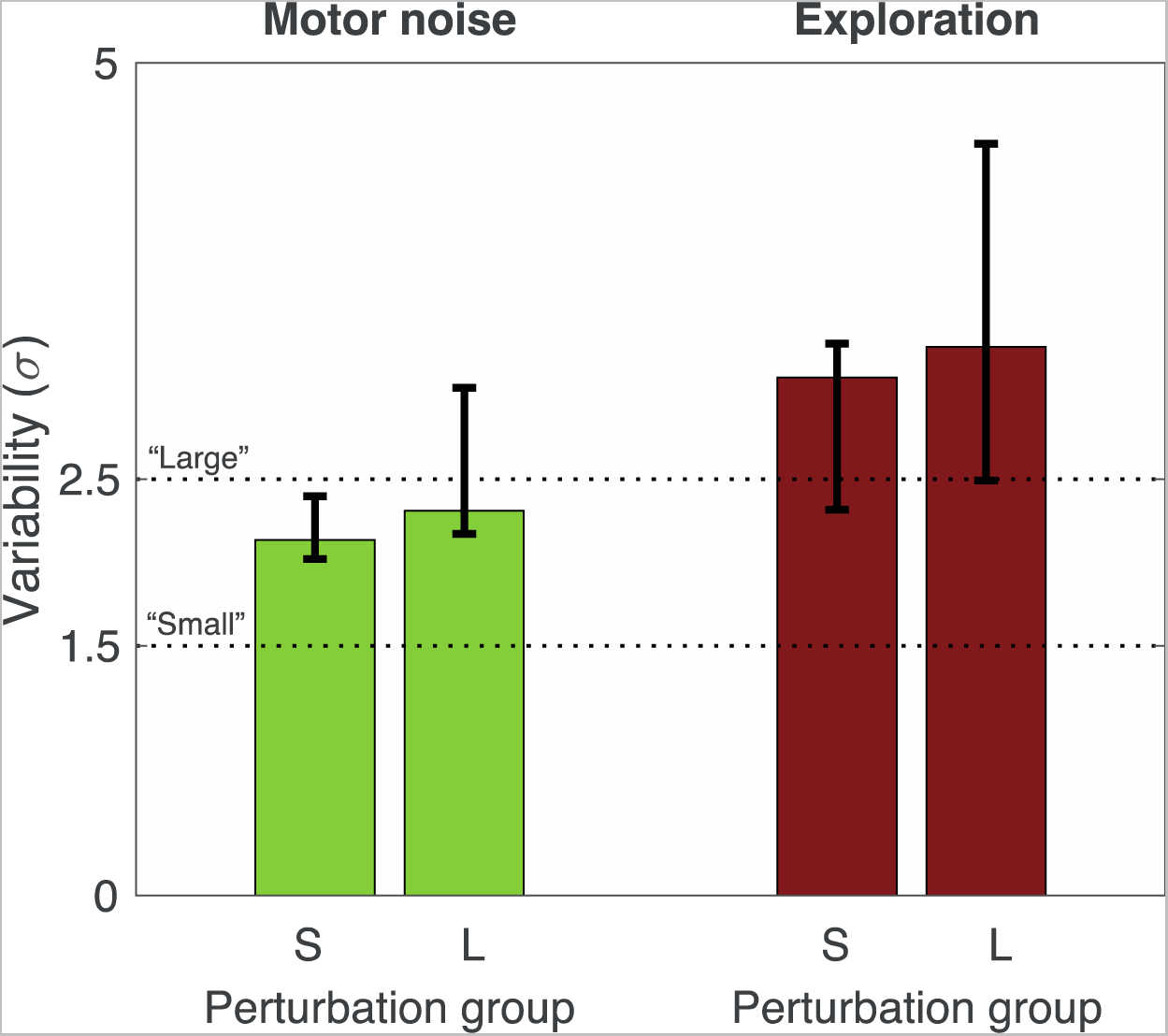
Reach angle variability separated into motor noise and exploration. Motor noise and exploration estimates are based on the ATTC-method with the simplest reward-based motor learning model (van Mastrigt et al., 2021). Median and interquartile range over participants for the Small perturbation group (S) and Large perturbation group (L). Horizontal dotted lines indicate step sizes of the gradual perturbation.

