## Supplementary material for "Implicit reward-based motor learning": Online resource 1. Questionnaires

---

**Experimental Brain Research**
Supplementary information

**Online resource 1** – Post-experiment questionnaires

**Post-experiment questionnaire**

Participant number:

Date:

Thanks for your participation! We have some questions for you!

The experimenter told you about two experimental groups:

- For the Aligned Group, the cursor position was always in alignment with the hand position.
- For the Misaligned Group, the cursor position was slightly displaced from the hand position.

1. Please indicate which group you were in:

   Aligned: _________

   Misaligned: _________

2. Please indicate your confidence in your judgment about your group assignment:

| 1 | 2 | 3 | 4 | 5 | 6 | 7 |
|---|---|---|---|---|---|---|
| No confidence (guessing) | | | Moderately Confident | | | Very Confident (sure) |

3. We want to know where you aimed relative to the center of the blue, middle target when you were doing the task and hearing the bings and buzzes.

Where did you aim when reaching to the blue, middle target?

Relative to the center of the blue, middle target, I aimed:

- To the left.
- To the center.
- To the right.

4. We now want to know where you aimed when doing the task in the last phase of the experiment when you were only hearing knocks and had to aim to different targets.

   Where did you aim when reaching to the blue, middle target?

   Relative to the center of the blue, middle target, I aimed:

   - To the left.
   - To the center.
   - To the right.

5. You were actually in the Misaligned Group, the one in which the cursor was slightly displaced from the position of your hand. Do you think the cursor was displaced to the left of your hand or displaced to the right of your hand?"

Version 1 (used for the Small perturbation group):

- Cursor was displaced to the left, as in the picture below.

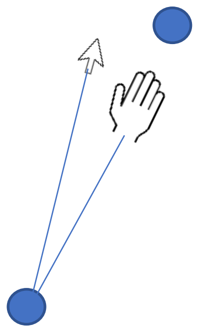

- Cursor was displaced to the right, as in the picture below.

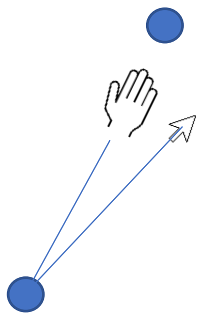

Version 2 (used for the Large perturbation group):

- Cursor was displaced to the left, as in the picture below.

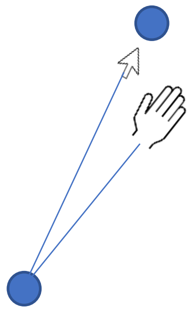

- Cursor was displaced to the right, as in the picture below.

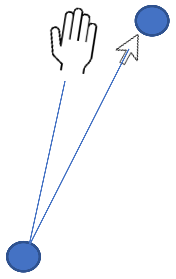
