## Supplementary material for "Implicit reward-based motor learning": Online resource 2. Questionnaire and behavioural results

---

**Experimental Brain Research**
Supplementary information

**Online resource 2** – Post-experiment questionnaire results

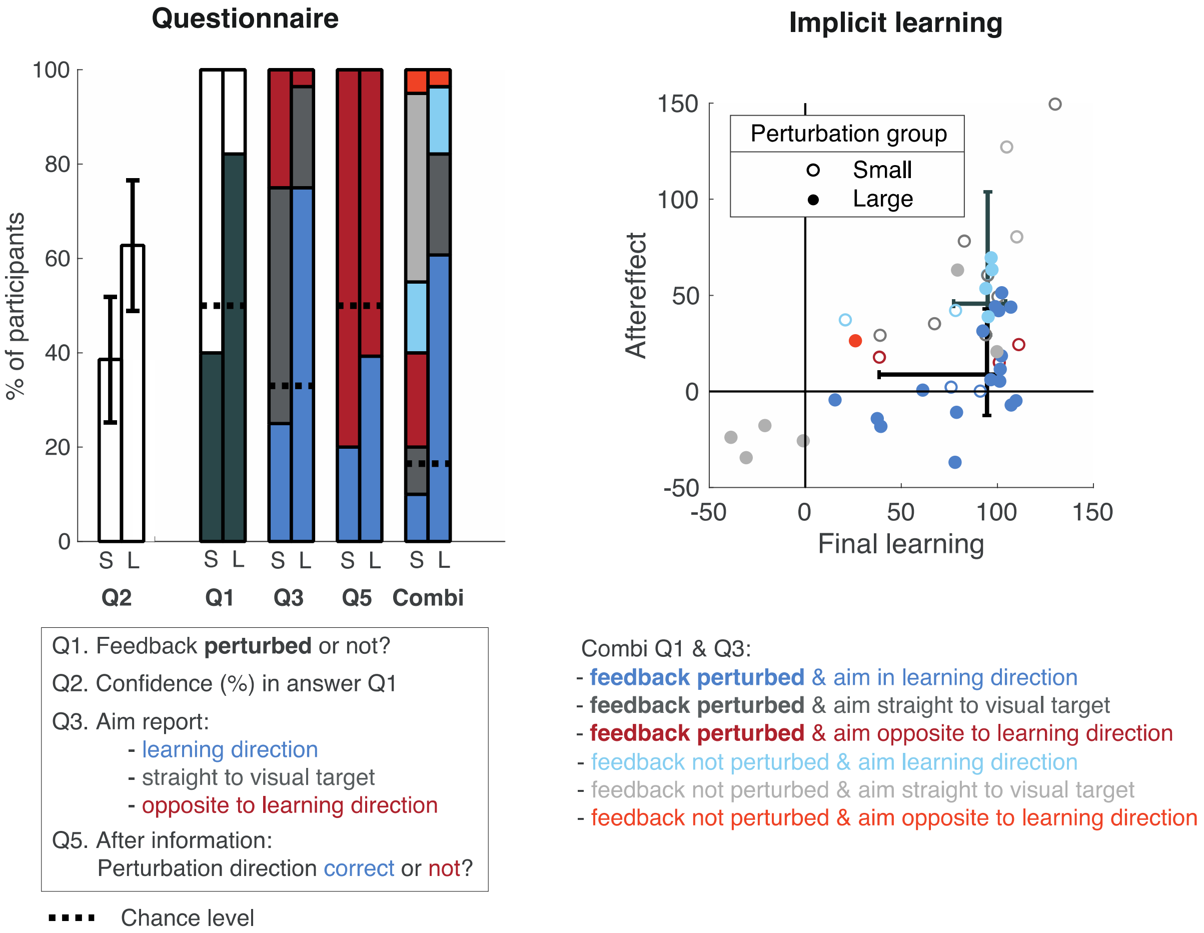


*Online resource 2. Questionnaire results related to learning for the Small Perturbation group (left bars) and the Large Perturbation group (right bars). Q's correspond to question numbers on the questionnaire. See Online resource 1 for the post-experiment questionnaires for the Small perturbation group and Large perturbation group.*
