## Supplementary material for "Implicit reward-based motor learning": Online resource 3. Learners and non-learners

---

**Experimental Brain Research**
Supplementary information

**Online resource 3** – Learners and non-learners

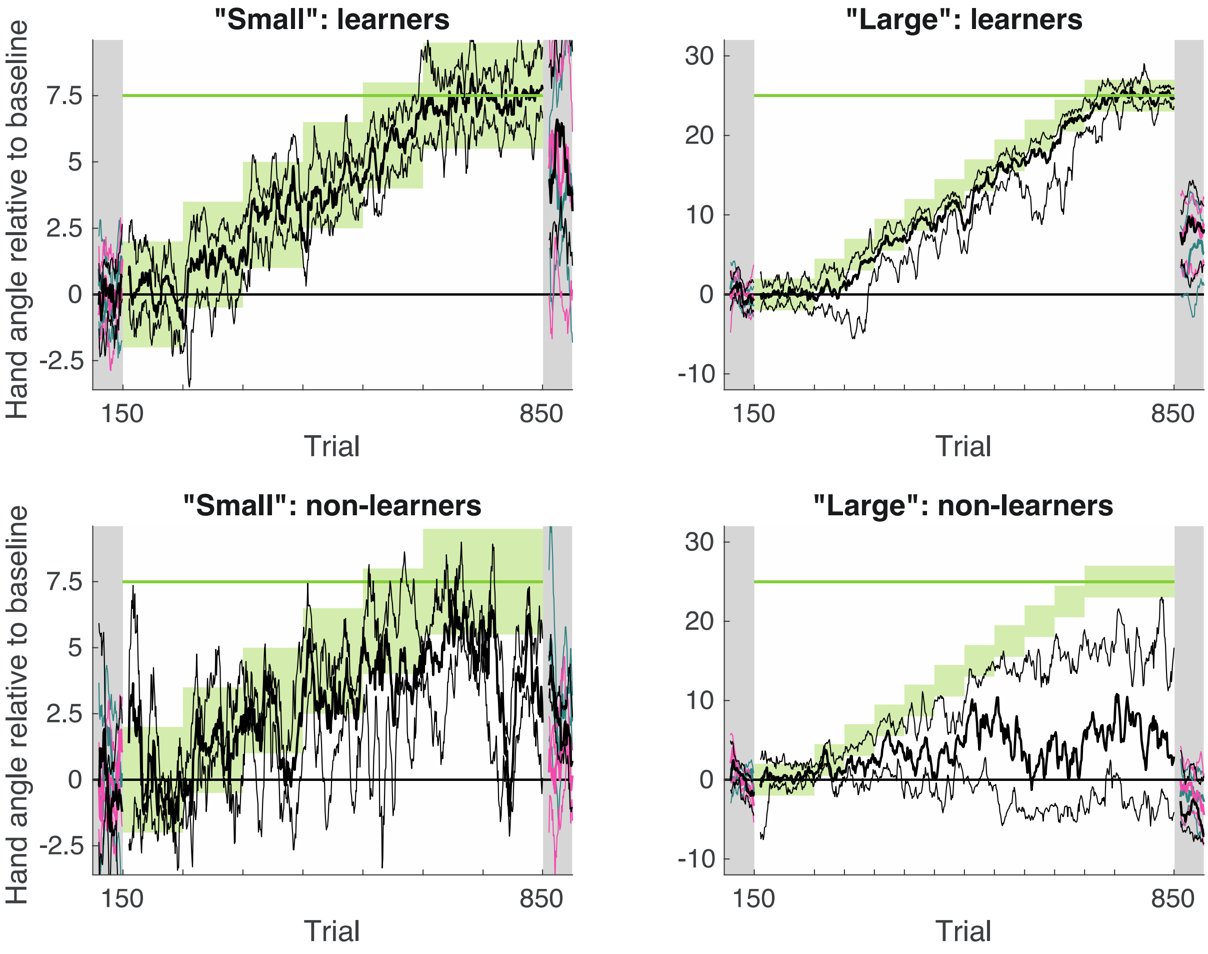


*Online resource 3. Learners and non-learners. Participants were divided into two groups based on whether final learning was in the reward zone ("learners", top rows) or not ("non-learners", bottom rows). By this definition, there were 16 learners and 4 non-learners in the Small Perturbation group (left) and 16 learners and 12 non-learners in the Large Perturbation group.*
