## Supplementary material for "Implicit reward-based motor learning": Online resource 4. Reach angle variability

---

**Experimental Brain Research**
Supplementary information

**Online resource 4** – Reach angle variability

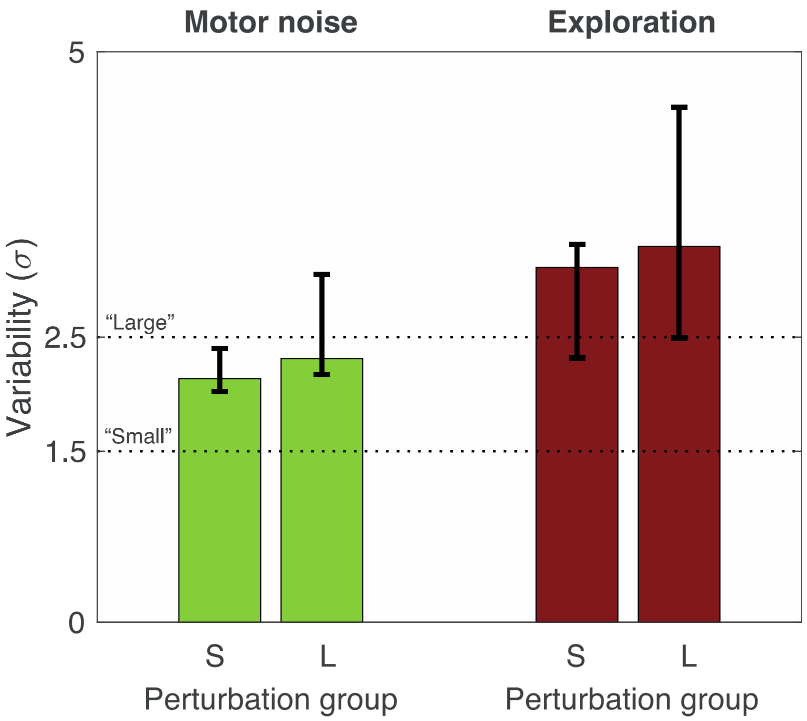


*Online resource 4. Reach angle variability separated into motor noise and exploration. Motor noise and exploration estimates are based on the ATTC-method with the simplest reward-based motor learning model* (van Mastrigt et al., 2021). *Median and interquartile range over participants for the Small perturbation group (S) and Large perturbation group (L). Horizontal dotted lines indicate step sizes of the gradual perturbation.*
